# Dissecting serum polyclonal antibody escape to SARS-CoV-2 variants by deep mutational learning

**DOI:** 10.1101/2025.05.01.651552

**Authors:** Danielle Shlesinger, Viktor Sadilek, Mason Minot, Evangelos Stamkopoulos, Thomas Bikias, Raphael Kuhn, Andreas Agrafiotis, Joseph M. Taft, Alexander Yermanos, Sai T. Reddy

## Abstract

The rapid emergence of SARS-CoV-2 variants harboring multiple receptor-binding domain (RBD) mutations continues to challenge the efficacy of vaccines and antibody therapeutics. While deep mutational scanning (DMS) has been instrumental in mapping single-mutation effects on antibody binding and immune escape, it remains limited in its ability to assess combinatorial mutational landscapes. Here, we extend deep mutational learning (DML), a method integrating combinatorial mutagenesis, yeast surface display, deep sequencing, and machine learning, to analyze serum polyclonal antibody escape. Human sera from COVID-19-vaccinated individuals were screened against diverse RBD variant libraries, validating 300 serum-variant interactions across 10 individuals by comparing predicted and observed binding to 30 RBD variants, confirming accurate mapping of binding and escape profiles. Model performance remained consistent across machine learning architectures, suggesting that serum binding and escape are governed by distinct, localized RBD sequence features. Notably, escape profiles were highly individualized, whereas binding signatures were more conserved, reflecting convergent epitope targeting. This approach highlights the potential of DML to generalize beyond observed RBD variants, assess cohort-specific immune breadth, and inform vaccine and therapeutic design in the face of viral evolution.

## Introduction

Since the start of the SARS-CoV-2 pandemic, the emergence of highly transmissible and immune-evasive variants has limited vaccine efficacy and driven waves of infection, posing challenges to sustained immunity at the population level (Tao et al. 2021; Eguia et al. 2021). The receptor-binding domain (RBD) of the Spike protein facilitates viral entry by specifically interacting with the angiotensin-converting enzyme 2 (ACE2) receptor on host cells. Spanning amino acids 333–527, the RBD is the target of approximately 90% of neutralizing antibodies (Piccoli et al. 2020). Mutations in the RBD can lead to an increase in binding affinity to ACE2 and also reduce the effectiveness of neutralizing antibodies, thereby facilitating immune escape (Ma et al. 2023). Traditional methods for studying antibody responses, such as serum/plasma titers by enzyme-linked immunosorbent assays (ELISA) are limited in throughput in regards to the number of samples and variants screened. Given the importance and abundance of mutations in the RBD that contribute to escape from neutralizing antibodies, more advanced, multidimensional approaches are needed to analyze how the mutational sequence landscape correlates with immune evasion (Cao et al. 2022; Lopez-Morales et al. 2023; Greaney, Loes, et al. 2021).

The protein engineering method of deep mutational scanning (DMS) (Fowler & Fields 2014) coupled to yeast surface display screening has been used extensively to investigate how single-position mutations in the RBD affect binding to ACE2 and neutralizing antibodies, thus revealing residues and mutations that drive immune escape (Starr et al. 2020; Starr, Greaney, Dingens, et al. 2021; Greaney, Loes, et al. 2021; Starr, Greaney, Addetia, et al. 2021; Starr, Czudnochowski, et al. 2021; Greaney, Starr, Barnes, et al. 2021; Greaney, Starr, Gilchuk, et al. 2021; Starr et al. 2022; Cao et al. 2022; Cao et al. 2023). Additionally, screening DMS libraries against human plasma has facilitated comprehensive mapping of RBD mutations that reduce binding to polyclonal antibodies (Greaney, Loes, et al. 2021). While DMS can yield variants containing multiple mutations, these variants arise stochastically and represent only a small fraction of the combinatorial mutational space.

The emergence of Omicron and its widely circulating sublineages, with more than 15 RBD mutations compared to the ancestral SARS-CoV-2 (Wu-Hu-1), have highlighted the need to assess combinatorial mutations; however, the vast combinatorial mutational sequence space of the RBD exceeds current experimental capabilities (Cao et al. 2022; Taft et al. 2022). To address these limitations, we have established the method of deep mutational learning (DML), which consists of screening combinatorial mutagenesis libraries of the RBD for binding or escape to ACE2 and neutralizing monoclonal antibodies (mAbs), followed by deep sequencing; this data is used to train supervised machine learning (ML) classification models that can accurately predict if an RBD variant binds or escapes ACE2 or neutralizing antibodies based on the sequence of the RBD (Taft et al. 2022). DML has also been used to determine the breadth of antibodies, allowing for the selection of neutralizing mAb combinations that optimally cover the RBD mutational sequence space (Frei et al. 2025) and to investigate adaptive mutational trajectories by synthetic co-evolution (Ehling et al. 2024).

The application of DML to serum polyclonal antibodies offers an unexplored but promising approach to study immune evasion driven by combinatorial mutations, particularly in profiling how existing or synthetic (hypothetical) viral variants escape polyclonal immune responses. Here, we apply DML to screen human serum antibody reactivity from individuals post-COVID-19 vaccination against combinatorial yeast-displayed SARS-CoV-2 RBD libraries, integrating deep sequencing and ML to predict variant binding and immune escape. Our ML models accurately predicted RBD escape mutants, were validated experimentally, and demonstrated generalizability to unseen sequences, while also revealing how the experimental screening workflow and ML model architectures critically influence predictive outcomes. By determining serum polyclonal antibody escape fingerprints, this approach advances variant risk assessment and can serve as an effective immune-monitoring tool to discern and evaluate the protective breadth of current and future vaccines.

## Results

### Serum profiling of SARS-CoV-2 RBD variants reveals patterns of binding and immune escape across individuals

DML has thus far only been applied to study SARS-CoV-2 RBD escape profiles in the context of neutralizing mAbs (Taft et al. 2022; Frei et al. 2023). We therefore adapted DML to predict RBD variant escape from polyclonal serum antibodies obtained from COVID-19 vaccinated individuals. To accomplish this, we leveraged three previously published SARS-CoV-2 RBD mutagenesis libraries, referred to as 1C, 2C, and 3C, which were designed using the ancestral Wu-Hu-1 sequence as a reference, and correspond to different epitope sites of the RBD, referred to as receptor binding motifs (RBM) (RBM-1 [residues 452-478], RBM-2 [residues 484-505] and RBM-3 [residues 440-452]). These libraries, constructed using combinatorial mutagenesis informed by deep mutational scanning (DMS) data, were designed to cover a high mutational sequence space with theoretical amino acid diversities ranging from 3.5×10^7^-1.5×10^10^ and were previously isolated based on binding to human ACE2 receptor (Taft et al. 2022). To focus on biologically relevant RBD variants that show functional receptor interactions, we pooled the ACE2+ fractions of the three separately assembled 1C, 2C and 3C libraries in equal ratios into one combined library. We screened this library against ten serum samples obtained from individuals following two-dose vaccinations (6-18 days post-second vaccination) during the early states of the pandemic (early 2021) (Table S1). The serum was heat-inactivated and pre-cleared with uninduced yeast cells to remove yeast binding antibodies prior to fluorescence-activated cell sorting (FACS). Serum binding and escape variants were isolated and targeted deep sequencing (Illumina) of the RBD region was performed (Figure 1A, B and Figure S1A, B). In total, 3,037,678 binding and 1,423,537 escape sequences were retrieved, with overall fewer sequences originating from sublibrary 1C compared to 2C and 3C (Figure 1C, Figure S1C).

**Figure 1.**
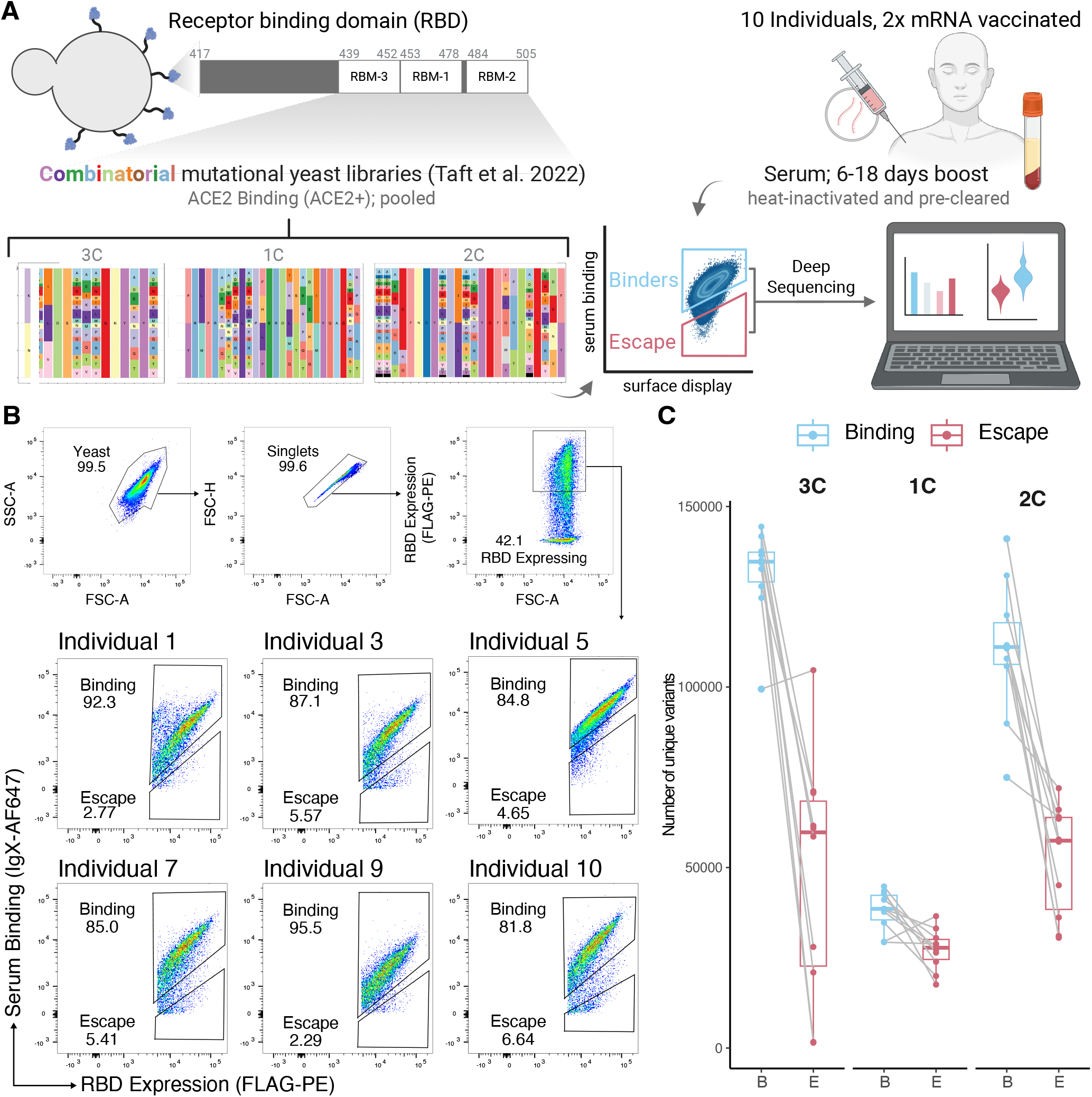
Overview of the screening of polyclonal serum against combinatorial yeast display libraries of SARS-CoV-2 RBD variants. A) Workflow illustrating the screening of polyclonal serum followed by deep sequencing and bioinformatic analysis. Yeast surface display library with combinatorial mutations designed in the RBMs of the RBD was screened against heat-inactivated pre-cleared serum by FACS. Binding and escape populations were isolated and subsequently deep sequenced. Binding separability was evaluated and validated, ensuring suitability for downstream training of machine learning models to predict serum-binding and -escape. B) Example flow cytometry dot plots depicting screening of ACE2+ 1C, 2C and 3C pooled library against serum. C) Paired number of unique serum-binding or -escape RBD variants retrieved from deep sequencing for each individual in each library.

We first examined the extent of sequence overlap between the classification labels of binding and escape within and across individuals, as the presence of overlapping RBD sequences (mixed labels) could complicate the training of downstream ML models. To this end, we calculated the Jaccard index (defined as the intersection divided by the union) within and across individuals, which demonstrated that binding and escape fractions showed varying degrees of similarity within and across individuals. Notably, serum antibody-binding RBD variants were significantly more shared across individuals compared to serum antibody-escape variants (Figure 2A). We further examined whether removing sequences with mixed labels within or across individuals would impact different similarity measures. Filtering decreased similarities overall, with binding sequences remaining more similar to each other than escape variants. Protein sequence logo plots revealed high similarity in amino acid usage between serum-binding and -escape variants, which was only minimally affected by filtering strategy (Figure 2B, S1B, S1C). As SARS-CoV-2 variants mutate to evade immune protection, we hypothesized that the sorted RBD escape variants would demonstrate larger mutational distances to the ancestral Wu-Hu-1 sequence compared to serum-binding sequences. We therefore calculated the Levenshtein distance in both serum-binding and -escape sequences across all RBMs. Serum-escape variants demonstrated a slightly higher Levenshtein distance from the Wu-Hu-1 across all RBMs. These differences were more pronounced when mixed-label sequences were removed, highlighting that removing mixed-label sequences may improve class separability (Figure 2C, S1B, S1C). Given the sequence similarity of serum-binding and -escape variants and their FACS gating proximity, we next experimentally validated the binding properties of observed sequences. To this end, we selected eight RBD variants that were present exclusively in binding or escape fractions in all ten individuals, produced them as single-variants by yeast surface display and tested them for serum binding by flow cytometry (Table S2). We calculated the normalized serum reactivity, which is defined as the mean ratio of the serum binding signal (anti-IgX) over the RBD-expression signal (anti-FLAG) for each variant and compared this to the mean ratio of the Wu-Hu-1. The calculated ratio reflects the normalized reactivity of the serum against the RBD, accounting for variations in RBD expression. In contrast to the observed binders, observed escape variants had significantly lower normalized serum reactivities compared to Wu-Hu-1. Directly comparing observed serum-binding and -escape variants demonstrated significantly different normalized serum reactivities, further validating the sensitivity of our yeast display and FACS workflow to produce accurate labels (Figure 2D, S1C). Although sequence similarity exists between serum-binding and -escape RBD variants, sorting combinatorial mutagenesis yeast libraries against polyclonal serum once can recover millions of RBD sequences, with serum-escape variants that exhibit higher mutational distances than binders.

**Figure 2.**
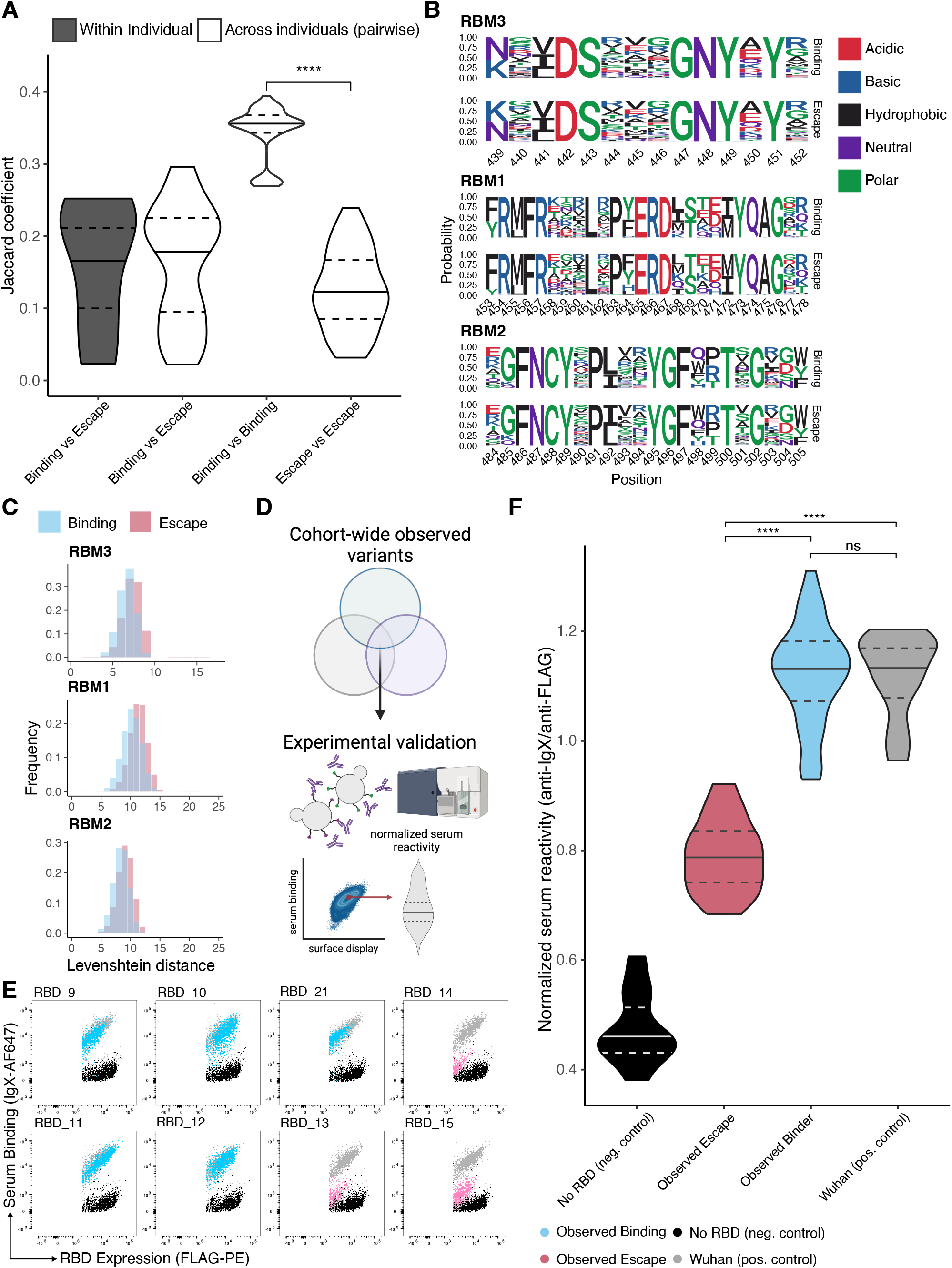
Screening of polyclonal serum against combinatorial yeast display libraries can recover separable serum-binding and -escape RBD variants with experimentally verified binding profiles. A) Violin plot depicting Jaccard index between binding and escape fractions within and across individuals. B) RBM protein sequence logo plots from serum-binding and -escape variants. Sequences with mixed labels were removed across the cohort. C) Comparison of distributions of mutational distances of RBMs from serum-binding and -escape fractions. Mutational distance was measured as Levenshtein distance of an RBM from the Wu-Hu-1 sequence. Sequences with mixed labels were removed across the cohort. D) Overview of validation experiments for observed variants. Cohort-wide binding and escape variants were expressed individually on yeast cells and screened against all individual serum samples by flow cytometry. Normalized serum reactivity was calculated by extracting the mean ratio of the serum binding signal (anti-IgX) over the RBD-expression signal (anti-FLAG) E) Representative FACS plots from screening one serum against observed RBD variants. F) Group comparison of normalized serum reactivity (mean signal ratio anti-IgX/anti-Flag) for observed serum-binding and -escape variants. Statistical analysis was performed using the unpaired two-tailed Student’s t test. ∗p<0.05, ∗∗p<0.01, ∗∗∗p<0.001, ∗∗∗∗p<0.0001, not significant (ns).

### Individual specific and cohort-wide classifiers accurately predict serum-escape variants

DML relies on using ML models to accurately predict binding and escape profiles of experimentally unscreened RBD variants (Taft et al. 2022; Frei et al. 2025), thereby enabling extrapolation over a broader mutational sequence space compared to experimental methods such as DMS. Mixed labels can introduce ambiguity and impair a classification task regardless of whether a mixed label accurately represents the underlying data. Our previous results uncovered the presence of mixed labels both within and across individuals (Figure 2A). To address ambiguity in RBD variants appearing in both binding and escape groups, we employed two complementary strategies: (i) Individual-specific filtering, removing sequences with conflicting labels within each donor and training 10 donor-specific classifiers to capture personalized RBD serum profiles (Figure 3A); and (ii) Cohort-wide filtering, eliminating all overlapping sequences across the entire cohort to train a single consensus model. This dual approach distinguishes whether mixed labels reflect biological individual variation (supported by superior individual model performance) or technical noise (indicated by cohort model performance) (Figure 3A).

**Figure 3.**
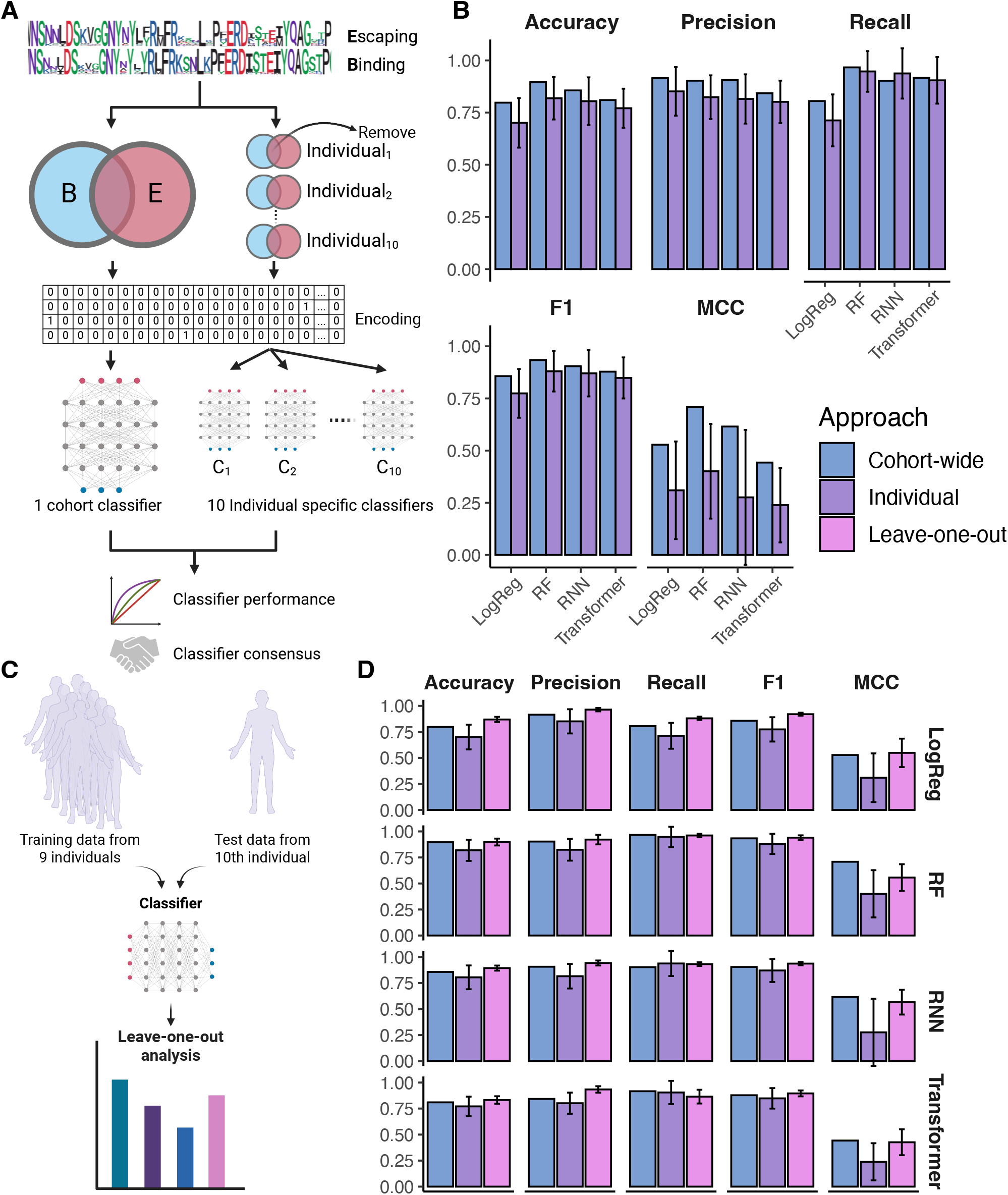
Cohort-wide machine learning models accurately predict serum antibody-binding and escape-variants. A) Computational workflow for the machine learning analysis of serum screened RBD variants: Recovered sequences were either stratified into cohort-level (left) or individual-specific (right) sets of serum-binding and -escape variants. Variants with mixed class labels were removed from the data before either individual-specific or cohort-wide classifiers were trained. Classifiers were trained to predict the serum binding or escape phenotype of an RBD based on sequence. B) Comparison of performance metrics across averages of different individual-specific classifiers or a cohort classifier (logistic model [LM], random forest [RF], recurrent neural network [RNN]), Transformer. C) Computational workflow of the “leave-one-out” approach. D) Ten RF classifiers were trained using a ‘leave one out’ strategy leaving the last individual out for classifier performance evaluation. Comparison shown to individual-specific classifiers and cohort classifiers. Standard deviations are shown by error bars.

In our previous DML study, we utilized an ensemble model approach with random forests (RF) and recurrent neural networks (RNN) (Taft et al. 2022). To capture both position-specific and longer-range sequence-function relationships, here we additionally included a logistic regression model for interpretable parameter-low predictions and a protein language model (PLM) embeddings, ESM2-650M (Transformer-based), paired with logistic regression classification head to incorporate and deep contextual learning via transfer learning for parameter-rich modeling (Bikias et al. 2025). We screened serum against the pooled 1C, 2C and 3C sub-libraries and subsequently trained all ML models on the combined RBD sequence region (position 417-505). To reduce bias in the training data we balanced both for label and sub-library, ensuring an equal representation of serum-binding and -escape variants, as well as variants from each sub-library. In both approaches and across all tested architectures, the RF model best predicted RBD escape-variants, with cohort-wide classifiers achieving F1 and MCC scores of 0.93 and 0.71, respectively, whereas individual classifiers resulted in mean values of 0.88 and 0.40, respectively (Figure 3B). The performance of individual classifiers varied strongly across individuals, and correlated with the number of sequences (amount of training data) retrieved for the corresponding individual (Figure S3A). This correlation underscores the benefit of the cohort-wide classifier, as the cost of deep sequencing escalates with increasing cohort sizes. We next assessed the ability of the cohort-wide classifier to generalize to unseen individuals and determined whether individual-specific patterns might limit its effectiveness. To this end we performed a ‘leave-one-out’ training and testing approach, corresponding to ten different cohort-wide classifiers (Figure 3C), each of which was trained using data from all individuals except one. To ensure there was no data leakage between training and test data, any overlapping sequences observed in a given individual and the rest of the training data were filtered out. This approach was comparable to the cohort-wide approach and resulted in a mean F1 score of 0.94 and MCC of 0.55 (Figure 3D). This implies that the distribution of serum antibody binding- and escape-variants learned at the cohort level is reflected at the individual specific level. At the resolution of yeast display and the available sequencing depth, individual antibody landscapes appear broadly similar, with no detectable differences in individual escape patterns (Figure 4S). Thus, cohort-level classifiers may perform as effectively as individual-specific models in predicting serum antibody binding and escape profiles.

**Figure 4.**
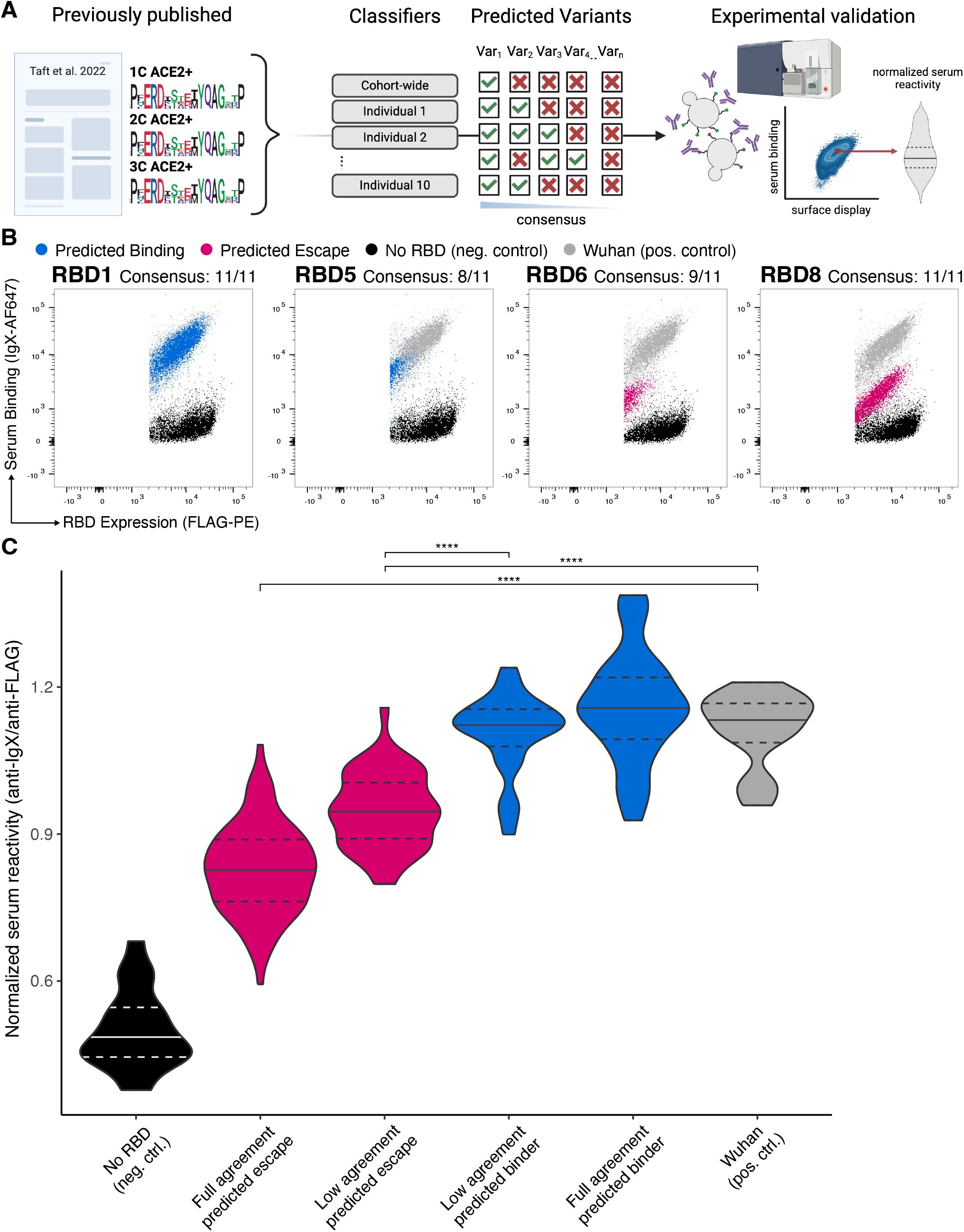
High consensus predictions of classifiers trained are experimentally validated. A) Workflow for the experimental validation of unseen RBD variants (not present in training or test data). B) Representative FACS plots from screening serum against unseen RBD variants, where the ensemble of all models had full agreement. C) Comparison of normalized serum reactivities (anti-IgX/anti-Flag) for predicted serum binding- and -escape variants with either full or low ensemble model agreement. Statistical analysis was performed using the unpaired two-tailed Student’s t test. ∗p<0.05, ∗∗p<0.01, ∗∗∗p<0.001, ∗∗∗∗p<0.0001, not significant (ns).

Screening yeast libraries often requires multiple enrichment rounds to reduce noise, which can be labor-intensive, costly and limited by sample availability (Traxlmayr 2022). We conducted two enrichment rounds for four individuals, finding that while enrichment reduced overlap between binding and escape fractions (Jaccard indices: 0.001–0.085 within individuals, 0.001–0.082 across individuals), it also increased the difference in Levenshtein distances between serum binding- and escape-variants, and enhanced classifier performance (F1 score: 0.98, MCC: 0.92) (Figure S3B, S3C, S3D). To reduce experimental effort, we created a mixed dataset by combining first-sort serum binding-variants with third-sort escape-variants, achieving comparable performance (F1 score: 0.99, MCC: 0.97) to fully enriched data (Figure S3F). Our results suggest that enriching only the escape population may be sufficient to generate high-quality training data while conserving experimental resources. Given the strong performance from single-sort data, the remainder of this study focuses on these single-sort datasets to further explore these outcomes.

### Experimental validation demonstrates DML can accurately predict serum binding- and escape-variants

After observing high accuracy of the RF classifier on predicting binding versus escape, we aimed to experimentally validate the performance of both the individual and cohort-wide classifiers on unseen RBD variants. These variants were not present in our training or test sets and were sourced from the previously published ACE2+ unpooled 1C, 2C and 3C libraries (Taft et al. 2022). To validate our predictions, we expressed 15 RBD variants individually on yeast that corresponded to full prediction agreement across all eleven models (cohort-wide and ten individual classifiers) and experimentally tested them for serum binding (Figure 4A). Furthermore, we ensured variants were sampled from different sub-libraries and varying mutational distances from Wu-Hu-1 (Table S3). Notably, across all predictions, there was generally more consensus for the prediction of binders, with 48.92% of the predicted variants classified with full consensus as binders, compared to only 0.04% as escape (Figure S5A). To analyze the serum binding phenotypes of all produced variants across all ten individuals, we again measured the normalized serum reactivity for each variant and compared this to the mean ratio of the ancestral Wu-Hu-1 RBD. Variants predicted to escape with full consensus showed significantly lower normalized serum reactivity compared to the positive control (Wu-Hu-1), whereas full consensus predicted binders had similar or higher normalized serum reactivity compared to the positive control. Predicted serum binding- and escape-variants substantially differed in their normalized serum reactivity, highlighting their distinguishable binding profiles.

Given the lower performance of our individual classifiers compared to the cohort-wide classifiers, we next focused on assessing the binding profiles of RBD variants where predictive discrepancies between models occurred. We selected and screened seven variants where the ensemble models did not reach full agreement. The selected variants were expressed on yeast and then evaluated for serum binding, and their normalized serum reactivity was measured. Those RBDs in which the majority of models predicted binding had similar normalized serum reactivity compared to the positive control, while variants partially predicted to escape had significantly lower normalized serum reactivity (Figure 4C). This demonstrated that when using our ensemble approach for prediction, the majority of the ensemble correctly coincides with serum binding and escape. Notably, the normalized serum reactivity of low agreement variants were closer to each other compared to full agreement predicted variants, indicating that the level of agreement could be associated with reduced separability of the serum reactivity. (Figure 4C, S5B, C).

We further investigated the models that deviated from the majority consensus. In only one of the seven produced variants did the cohort-wide classifier assign the wrong label, which also matched the minority prediction among all models (Table S4; RBD7). More detailed analysis of the normalized serum reactivity showed that low agreement binders were correctly predicted by the majority of the ensemble, highlighting our previous finding that our models show greater agreement in identifying binders (Figure S5A, S6A). In contrast, for low-agreement escape variants, one of the two minority models correctly defined the ensemble (Figure S6A; RBD6, individual 8). Moreover, there were also cases where single individual responses did not align with the majority prediction, with single individual serum reactivities being close to the positive control (Figure S6A). Notably, visual inspection of the FACS plots showed that many of the produced variants with low agreement had low expression profiles (Figure S6B). This suggests that disagreement among the models can reflect a uniqueness of individual antibody landscapes in rare cases, and additionally points to a challenging region of the sequence space that is difficult to predict. Overall, our findings suggest that a cohort-wide classification approach is generally sufficient to accurately predict serum binding profiles, as individual serum landscapes appear functionally similar when assessed with yeast display at the given sequencing depth. However, disregarding individual labels can reduce classification accuracy, underscoring the advantage of an ensemble approach despite its higher computational cost. These results demonstrate that even with single-sort data, DML can effectively predict serum binding and escape using ensemble modeling.

## Discussion

Here, we extend the application of DML to predict serum polyclonal antibody binding and escape across SARS-CoV-2 RBD variants. While prior DMS studies have validated single-mutation escape variants by identifying their enrichment in circulating immune-evasive strains and linking them to reduced neutralization capacity (Greaney, Loes, et al. 2021), these approaches are limited in capturing the complexity of combinatorial mutations. In a previous study, we demonstrated that DML could accurately identify multi-site mutations driving escape from therapeutic monoclonal antibodies, validating 46 synthetic RBD variants across four antibodies with over 90% accuracy and a total of 178 experimental validations (Taft et al. 2022; Frei et al. 2025). Building on this foundation, we now adapt the DML framework to analyze more complex polyclonal sera, addressing key challenges such as label noise, limited sample availability, and the inherent heterogeneity of individual serum responses in high-throughput experimental workflows.

High-throughput screening methods inherently produce noisy data, and FACS-based yeast surface display screens, commonly employed for mapping binding interactions, often require multiple rounds of enrichment to clearly distinguish binding from escape populations (Glick et al. 2003; Traxlmayr 2022). This iterative process is both time-consuming and constrained by limited sample availability, particularly when attempting to isolate rare binding events. Although recent meta-learning approaches have shown promise in accelerating screening and enhancing model robustness, they still rely on a small, trusted meta-set, typically derived from multiple rounds of enrichment (Minot & Reddy 2024). In this study, we demonstrate that DML can accurately recover distinct binding labels using only once-sorted data by computationally filtering mixed-label populations. While additional enrichment modestly reduced label ambiguity, it had only a minor effect on overall model performance, suggesting that selective enrichment of underrepresented populations may be sufficient, with computational strategies effectively compensating for reduced experimental depth.

Our DML analysis, consistent with prior DMS studies (Greaney, Loes, et al. 2021; Lee et al. 2019), revealed significant person-to-person variability as it relates to RBD mutations on serum escape, with discrepancies between individual responses and consensus predictions occurring predominantly for escape variants. In contrast, serum-binding variants were more consistently predicted across individuals, likely reflecting convergent targeting of shared neutralizing and non-neutralizing epitopes (Barnes et al. 2020; Cao et al. 2022; Dejnirattisai et al. 2021; Yuan et al. 2021; Amanat et al. 2021; Lima et al. 2022; Chen et al. 2021; Yan et al. 2024). These findings underscore the personalized nature of escape responses and the relative predictability of binding patterns, particularly within the RBD. Recent evidence suggests that immune history shapes the specificity of serum neutralizing antibodies (Dadonaite et al. 2025), implying that model generalizability may improve when stratifying individuals by shared immune backgrounds. Supporting this, we observed only minor performance differences among logistic regression, random forest, RNN, and transformer models, suggesting that escape and binding features are often governed by localized, interpretable sequence patterns. In the context of polyclonal sera, however, simpler models may suffice due to the dominant influence of direct sequence features and the obscured impact of conditional or allosteric interactions present in more focused antibody systems, particularly when training with limited sequence context.

While neutralizing antibodies are essential for blocking viral entry, antigen-binding antibodies also contribute to immune defense through alternative mechanisms, such as Fc-mediated effector functions (Abebe & Dejenie 2023; Zhang et al. 2023). Notably, certain mutations can reduce antigen binding without significantly impairing neutralization, likely due to conformational differences between isolated and native RBD structures (Greaney, Loes, et al. 2021). Tools such as the DMS-based “escape calculator” have shown correlations with neutralization assays for polyclonal sera but depend on the assumption that monoclonal antibody activity sufficiently captures the complexity of polyclonal responses (Greaney, Starr & Bloom 2021). Our study specifically examines vaccinated individuals, who tend to exhibit higher levels of non-neutralizing antibodies compared to those with natural infection (Amanat et al. 2021). Moving forward, it will be important to assess how yeast display-based serum binding assays relate to real-world protection or susceptibility to emerging SARS-CoV-2 variants. By integrating epidemiological and clinical datasets, DML has the potential to predict inter-individual variation in antibody responses shaped by immune history. Such insights could inform the selection of broadly effective antibody therapies and contribute to the design of new vaccines. Ultimately, this approach may help identify serum binding profiles linked to durable immunity, thereby enhancing our capacity to tailor interventions for sustained population-level protection.

## Methods

### Human serum samples

Serum samples collected under approval by the ethics committee of Sterling IRB (8291-BZhang) and purchased from RayBiotech through Anawa Trading SA (CUST-BB-11062021). Serum was obtained 6-18 days post vaccination with Pfizer-BioNTech or Moderna vaccine. All serum samples were heat-inactivated at 56°C for 30 min.

### Serum pre-clearing

To remove yeast-binding serum antibodies, serum was mixed with washed EB100 cells in wash buffer (Dulbecco’s PBS+ 0.5% BSA + 0.1% Tween20 + 2 mM EDTA). The mixture was incubated for 1 h at 4 °C with agitation (600-700 RPM). Yeast cells were pelleted (8000 x g, 30s) and the supernatant was collected and stored at 4 °C until use. Pre-cleared serum was diluted with wash buffer to the desired concentration, such that the fluorescent signal from RBD binding was similar across all samples (Greaney, Loes, et al. 2021).

### Screening RBD for serum binding and escape

ACE2 binding fractions of the previously described sublibraries 1C, 2C and 3C were pooled in equal ratios to form the 123C ACE2+ RBD library (Taft et al. 2022). Cells were cultured in SD-UT medium (20 g/L glucose, 6.7 g/L yeast nitrogen base without amino acids, 5.4 g/L Na2HPO4, 8.6 g/L NaH2PO4·H2O and 5 g/L casamino acids) at 30 °C in a 250 rpm shaking incubator. Surface expression was induced in SG-UT medium (20 g/L galactose, 6.7 g/L yeast nitrogen base without amino acids, 5.4 g/L Na2HPO4, 8.6 g/L NaH2PO4·H2O and 5 g/L casamino acids) at 23 °C for 36-42 h in a 250 rpm shaking incubator (Boder & Wittrup 1997). For each screen approximately 1.5 x 10^7^ library cells were washed twice with 1 mL of wash buffer (8000x g, 30 s) before staining. Primary staining was carried out with diluted serum samples, followed by secondary staining with anti-IgX (Alexa Fluor® 647 AffiniPure Goat Anti-Human IgA + IgG + IgM (H+L), Jackson ImmunoResearch 109-605-064, 1:200 dilution), and tertiary staining with anti-Flag (PE-anti-DYKDDDDK Tag Antibody, Biolegend 637310, 1:200 dilution). Cells were washed twice in between stainings, and resuspended with wash buffer and kept on ice until FACS or flow cytometry. Cells were sorted by FACS (BD FACSAria Fusion) and flow cytometry analysis was performed on the BD Fortessa cytometer. Collected cells were pelleted and cultured in SD-UT media until desired OD.

### Deep sequencing of RBD libraries

Deep sequencing was performed as previously described (Taft et al. 2022). In short: Plasmid DNA encoding the RBD variants was isolated following the manufacturer’s instructions (Zymo D2004). The region of interest was amplified using custom primers. Illumina Nextera barcode sequences were added in a second PCR amplification step, allowing for multiplexed high-throughput sequencing runs. Library preparation was performed using using SPRIselect beads according to the manufacturer’s instructions (Beckman Coulter B23318). Populations were pooled at the desired ratios and sequenced using Illumina 2 x 250 PE protocols (NovaSeq instrument).

### Experimental validation of selected RBD variants

RBD variant inserts were cloned into a pYD1 drop out plasmid, with a sfGFP in place of the RBD, using the NEBridge® Golden Gate assembly kit according to manufacturer’s instructions (NEB #E1601). The assembled product was transformed into E. coli DH5α Mix & Go! Competent Cells competent cells (Zymo Research T3007) according to manufacturer’s instructions. sfGFP negative cell colonies were picked and cultured overnight. The pYD1 plasmids were purified using the Zymopure midiprep kit (Zymo Research, D4200). The plasmids were then transformed into EBY100 yeast using the Frozen-EZ Yeast Transformation kit (Zymo Research, T2001) according to the manufacturer’s instructions and plated on SD-UT agar. Single colonies were picked and cultured in SD-UT medium and subsequently screened using a BD Fortessa cytometer. Normalized serum reactivity was calculated by extracting the serum binding signal (anti-IgX) and RBD-expressing signal (anti-FLAG) for each cell of each variant and calculating the ratio between the two (anti-igX/anti-FLAG).

### Data processing

Data was processed as previously described (Taft et al. 2022): Reads were paired, quality trimmed and assembled using BBDuk on R. After extracting the regions of interest, sequences were translated using the R package Biostrings (v2.7.1). All sequences were filtered using a threshold of read counts >3. Moreover, sequences of incorrect length, sequences containing mutations outside of the library design and duplicate sequences were filtered out.

For machine learning analysis further filtering of sequences was carried out. For individual specific classifiers RBD sequences occurring in both the serum binding and serum-escape population within the set of sequences of the given individual were removed. For cohort-wide classifiers the individual labels were removed before the same filtering approach was applied.

### Similarity calculations

Jaccard coefficients were defined as the size of the intersection divided by the size of the union of the RBD sequences recovered from each sorted fraction. Levenshtein distances were calculated compared to Wu-Hu-1 (YP_009724390.1).

### Machine learning model training and evaluation

#### Preprocessing

All code was built in Python (3.11.6) and data was prepared using Numpy (v1.23.5) and pandas (v1.4.2).

#### Logistic regression (LG), Random forest (RF), Recurrent neural network (RNN)

The data was split into a training and testing set using a 80/20 split. As not all sub libraries were equally represented in the dataset we balanced our training data by either upsampling the minority label class (binding/escape) within each RBM sublibrary, in the case of LG and RF using scikit-learn (v1.5). The long short-term memory (LSTM) recurrent neural networks (RNN) models were built using Keras and Tensorflow (v2.14.2), and data was balanced by weighing the majority class (binding/escape) within each RBM. Hyperparameter search was performed by doing a grid search (Table S5).

#### Transformer

Data was balanced using the same approach as for the RNN. Feature extraction was performed using ESM2-650M and embeddings were pooled in the sequence dimension using the “mean” pooling parameter from the PLMFit package (Bikias et al. 2025). Embeddings were then fed into a logistic regression head. Hyperparameter tuning was performed according to the PLMFit package workflow.

### Data visualization

Experimental workflows were done using Biorender. All other graphics were generated in R, using the packages ggplot2 (v3.4.2), ggpubr (v0.6.0) and ggseqlogo (v0.1), ggprism (v1.0.5). The packages readr (v2.1.4), reshape2 (v1.4.4), tidyverse (v2.0.0) were additionally used for data import and transformation. Experimental overviews were created using BioRender.com and final figures were assembled with Adobe illustrator.

## Acknowledgements

We thank Dr. Christian Beisel, Mirjam Feldkamp, Elodie Burcklen, and Ina Nissen at the ETH Zurich D-BSSE Genomics Facility Basel for their support and assistance. We additionally acknowledge and thank Di Tacchio Mariangela, Cavallini Chiara, and Gumienny Aleksandra from the D-BSSE FACS facility for their support. Funding: This work was funded in part by ETH Zurich, Foundation Immune Engineering for Global Child and Adolescent Health and the Basel Research Center for Child Health.

## Author contributions

D.S., S.T.R., and A.Y. contributed to study design. D.S., V.S., R.K., A.A., and J.M.T. participated in the material preparation. D.S., V.S., M.M., E.S., and T.B. contributed to computational analyses and pipelines. D.S., A.Y., and S.T.R wrote the manuscript, with input from all other authors.

## Declaration of interest

S.T.R. is a co-founder and holds shares of Engimmune Therapeutics AG and Encelta and Fy Cappa Biologics. S.T.R. holds shares of Alloy Therapeutics. S.T.R. is on the scientific advisory board of Engimmune Therapeutics, Alloy Therapeutics, Encelta and Fy Cappa Biologics. S.T.R. is a member of the board of directors for Engimmune Therapeutics.

### Declaration of generative AI and AI-assisted technologies in the writing process

During the preparation of this work the author(s) used ChatGPT, Perplexity and spell check for language refinement and editing. After using this tool/service, the author(s) reviewed and edited the content as needed and take(s) full responsibility for the content of the published article.

## Supplementary Data

**Table S1.** Metadata for individual serum samples used in this study. Individual is the simplified identifier used in this study, sample ID is the unique identifier given by vendor (RayBiotech), serum dilution refers to the specific dilution level at which each experiment was conducted, days post second vaccination indicated the sample collection time point, sex and type of mRNA vaccine is also indicated.

| Individual | Sample ID | Sex | Days post 2nd vaccine | Vaccine Manufacturer | Serum Dilution |
| --- | --- | --- | --- | --- | --- |
| 1 | V974C8 | Female | 8 | Pfizer-BioNTech | 1 : 400 |
| 2 | V985C7 | Male | 7 | Moderna | 1 : 400 |
| 3 | V977C10 | Female | 10 | Pfizer-BioNTech | 1 : 800 |
| 4 | V981C | Male | 15 | Moderna | 1 : 400 |
| 5 | V979C8 | Female | 8 | Pfizer-BioNTech | 1 : 200 |
| 6 | V967C | Male | 17 | Moderna | 1 : 100 |
| 7 | V980C9 | Female | 9 | Pfizer-BioNTech | 1 : 400 |
| 8 | V989C | Male | 6 | Pfizer-BioNTech | 1 : 400 |
| 9 | V964C9 | Female | 9 | Pfizer-BioNTech | 1 : 400 |
| 10 | V952C | Male | 18 | Moderna | 1 : 400 |

**Figure S1.**
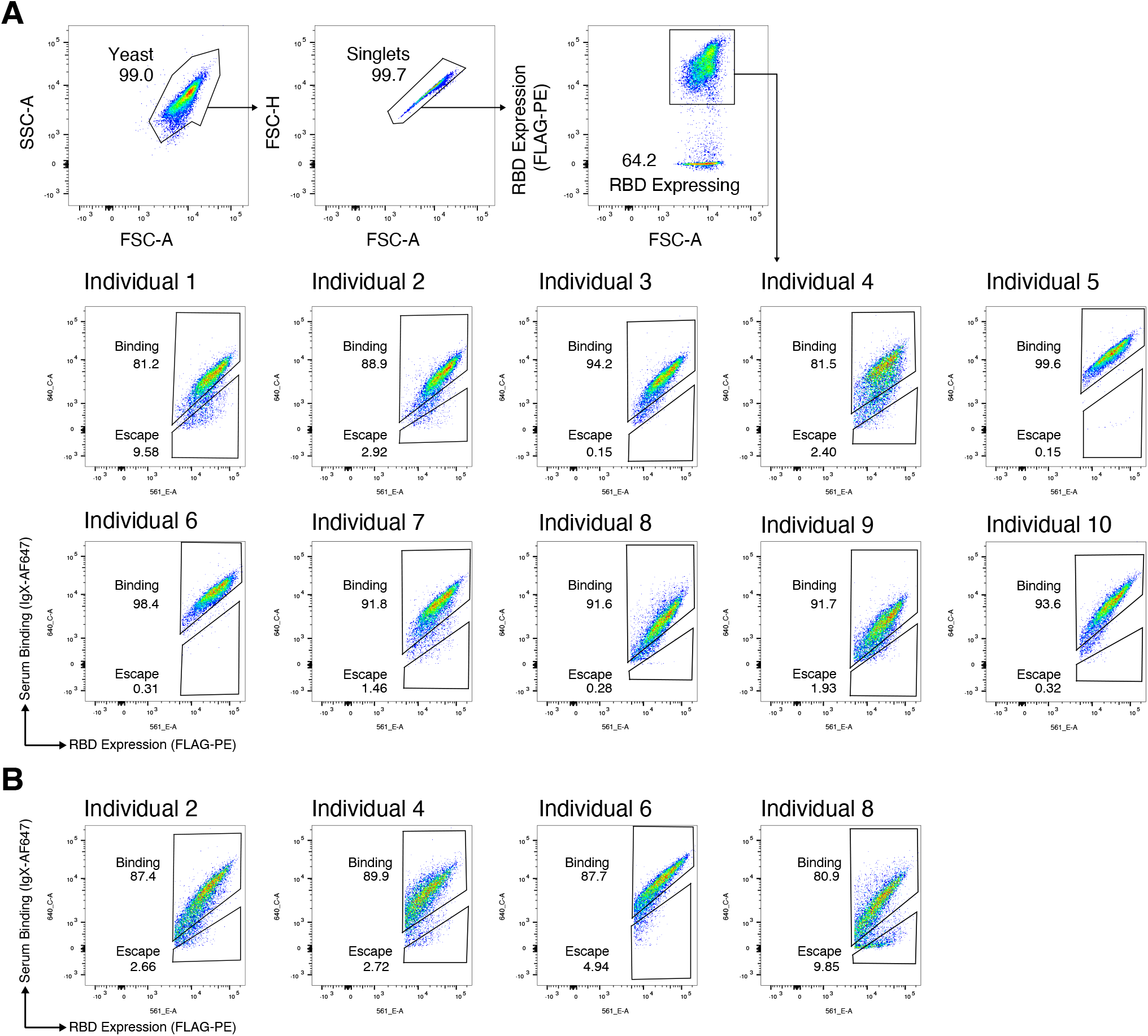
Yeast display screening of combinatorial RBD libraries for serum binding and escape. A) Control RBD (Wu-Hu-1) gating schemes used for selection of serum binding and -escape variants. B) Flow cytometry dot plots depicting screening of ACE2+ 123C library against serum of the individuals not included in Figure 1. Approximately 1.5 x 10^7^ yeast cells were screened for each individual serum.

**Figure S2.**
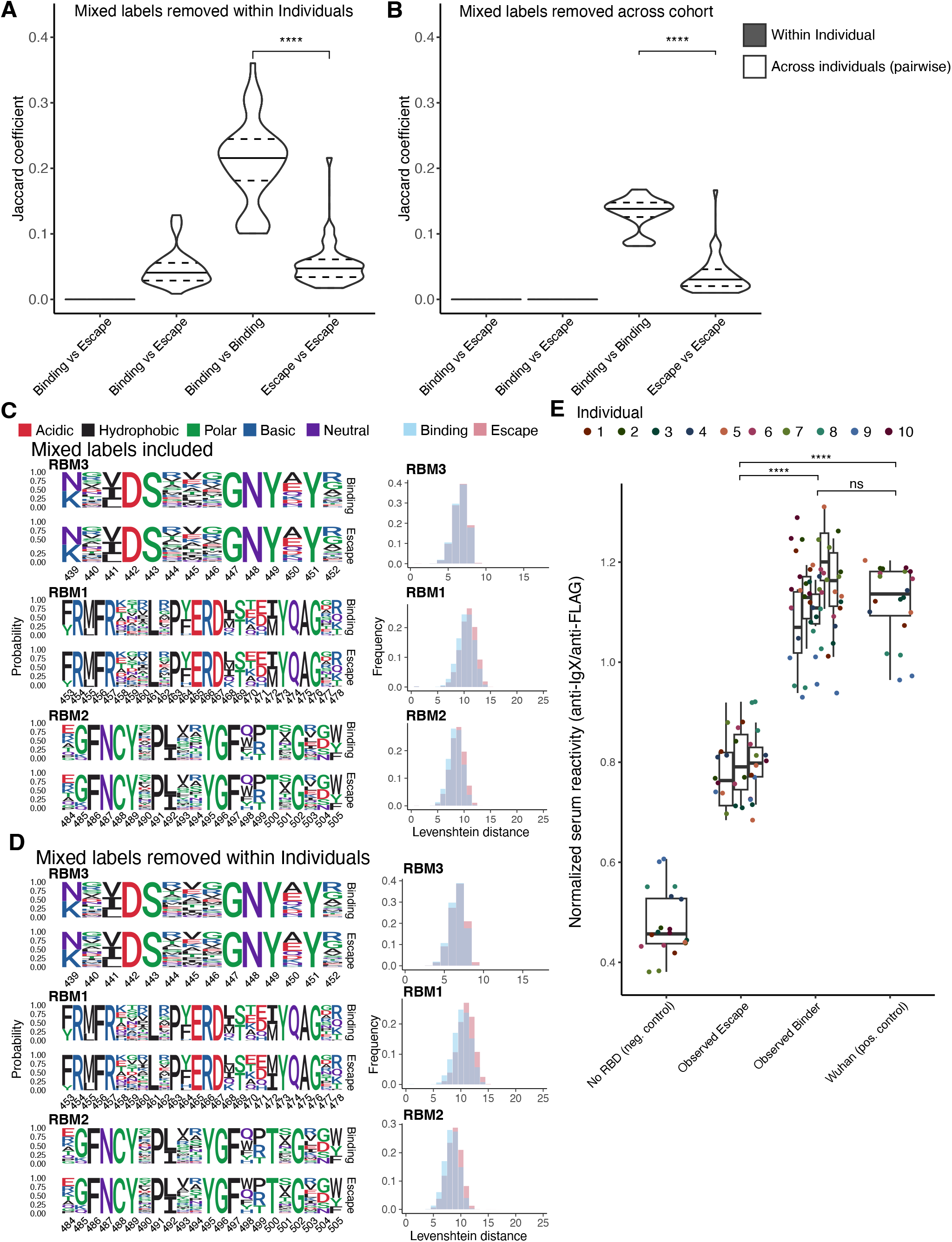
Additional characterization overlap, diversity and sequence similarity between serum binders and escape. A and B) Violin plot depicting Jaccard index between binding and escape fractions within and across individuals. Sequences with mixed labels were removed on an individual level (A) or cohort level (B). C and D) RBM sequence logo plots from serum-binding and -escape variants (left) and their comparison of distributions of mutational distances of RBMs and comparison of RBM diversity at each mutational distance (right). Sequences with mixed labels were included (C) or removed on an individual level (D). Mutational distance was measured as Levenshtein distance of an RBM from the Wu-Hu-1 sequence. E) Group comparison of normalized serum reactivity (mean signal ratio anti-IgX/anti-Flag ratios (anti-IgX/anti-Flag) for observed serum binding and -escape variants for all 8 tested observed variants. Statistical analysis was performed across groups using the unpaired two-tailed Student’s t test. ∗p<0.05, ∗∗p<0.01, ∗∗∗p<0.001, ∗∗∗∗p<0.0001, not significant (ns).

**Table S2.** Overview of validated observed RBD variants. Variant name, fraction from which it was retrieved (observed label), sub-library to which it belongs and Levenshtein distance from Wu-Hu-1 sequence.

| Variant | Observed label | Sub-library | Levenshtein distance | Sequence |
| --- | --- | --- | --- | --- |
| RBD_9 | Binding | 1C | 11 | KIADYNYKLPDDFTGCVIAWNSNNLDSKVGGNYNYLF<br>RMFRKARLRPYERDKTEEIYQAGAKPCNGVEGFNCYFP<br>LQSYGFQPTNGVGY |
| RBD_10 | Binding | 3C | 6 | KIADYNYKLPDDFTGCVIAWNSNEVDSQEGGNYDYVY<br>RLFRKSNLKPFERDISTEIYQAGSTPCNGVEGFNCYFPLQ<br>SYGFQPTNGVGY |
| RBD_11 | Binding | 3C | 6 | KIADYNYKLPDDFTGCVIAWNSNTIDSRPGGNYEYGYR<br>LFRKSNLKPFERDISTEIYQAGSTPCNGVEGFNCYFPLQS<br>YGFQPTNGVGY |
| RBD_12 | Binding | 3C | 4 | KIADYNYKLPDDFTGCVIAWNSNGVDSEVGGNYKYLY<br>RLFRKSNLKPFERDISTEIYQAGSTPCNGVEGFNCYFPLQ<br>SYGFQPTNGVGY |
| RBD_13 | Escape | 1C | 12 | KIADYNYKLPDDFTGCVIAWNSNNLDSKVGGNYNYLF<br>RMFRPRILVPLERDKTTEMYQAGNKPCNGVEGFNCYFP<br>LQSYGFQPTNGVGY |
| RBD_14 | Escape | 3C | 7 | KIADYNYKLPDDFTGCVIAWNSNTVDSVEMGNYKYPY<br>RLFRKSNLKPFERDISTEIYQAGSTPCNGVEGFNCYFPLQ<br>SYGFQPTNGVGY |
| RBD_15 | Escape | 3C | 6 | KIADYNYKLPDDFTGCVIAWNSNLIDSKIMGNYTYPYR<br>LFRKSNLKPFERDISTEIYQAGSTPCNGVEGFNCYFPLQS<br>YGFQPTNGVGY |
| RBD_21 | Binding | 2C | 8 | KIADYNYKLPDDFTGCVIAWNSNNLDSKVGGNYNYLY<br>RLFRKSNLKPFERDISTEIYQAGSTPCNGVEGFNCYRPIA<br>SYGFQRTYGQDF |

**Figure 3S.**
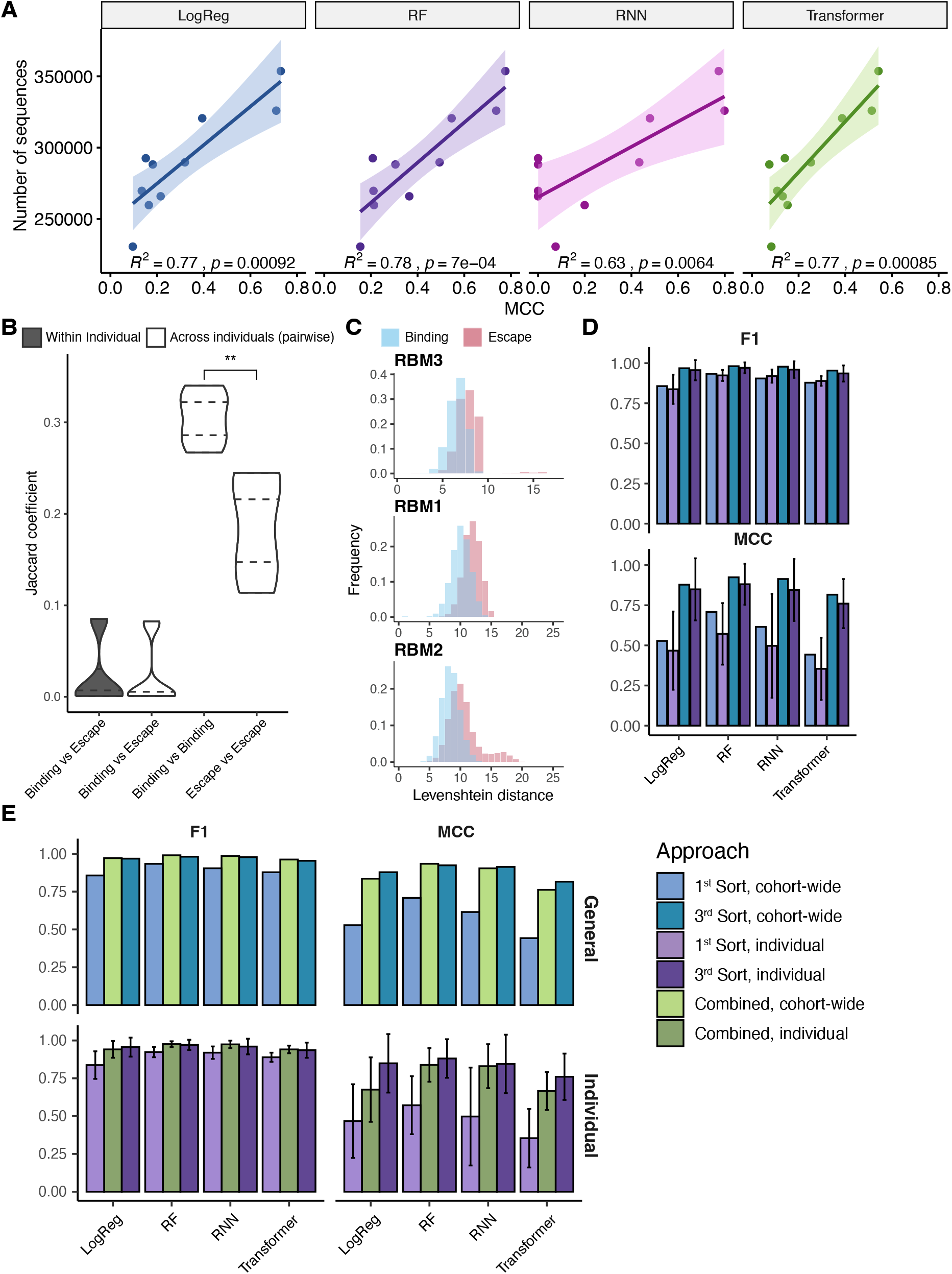
Correlation between training data, screening and machine learning model performance. A) Dot plots showing the correlation between MCC scores of individual classifiers and number of sequences retrieved for the corresponding individual. B) Violin plot depicting Jaccard coefficient between enriched binding and enriched escape fractions within and across individuals. C) Comparison of distributions of mutational distances of RBMs from serum binding and escape fractions and comparison of RBM diversity at each mutational distance of serum binding and escape variants. Mutational distance was measured as Levenshtein distance of an RBM from the Wu-Hu-1 sequence. D) Comparison of performance metrics across averages of different individual-specific classifiers or a cohort classifier (logistic model [LM], random forest [RF], recurrent neural network [RNN]), Transformer) trained on three times sorted and once-sorted data. E) Comparison of performance metrics across averages of different individual specific classifiers or a cohort classifier (logistic model [LM], random forest [RF], recurrent neural network [RNN]), Transformer) trained on once-sorted binding and three times sorted escape fractions. Statistical analysis was performed across groups using the unpaired two-tailed Student’s t test. ∗p<0.05, ∗∗p<0.01, ∗∗∗p<0.001, ∗∗∗∗p<0.0001, not significant (ns).

**Figure 4S.**
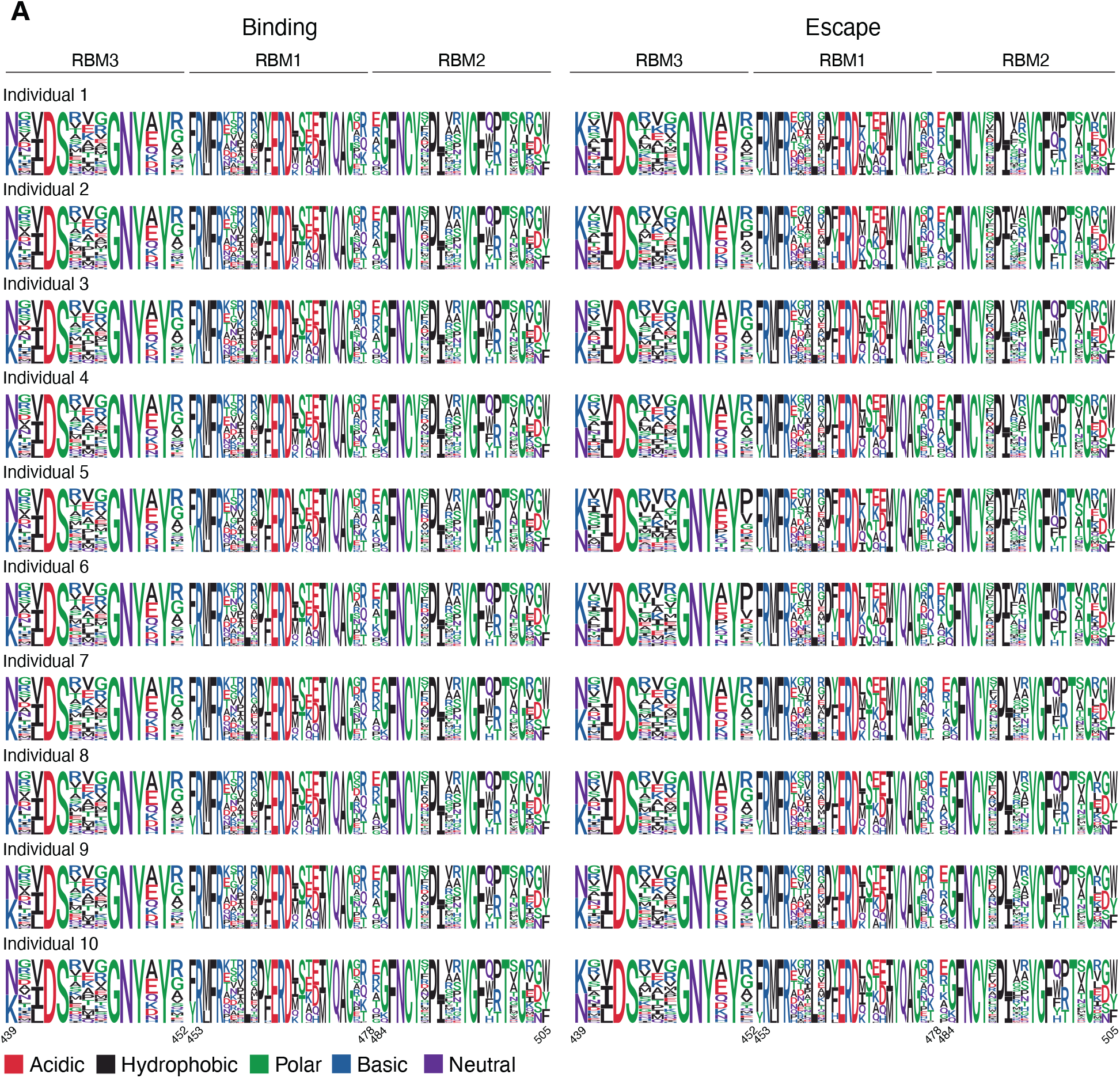
Individual RBM protein sequence logo plots from serum-binding and -escape variants. A) RBM sequence logo plots from serum-binding and -escape variants. Sequences with mixed labels were removed on an individual level.

**Figure S5.**
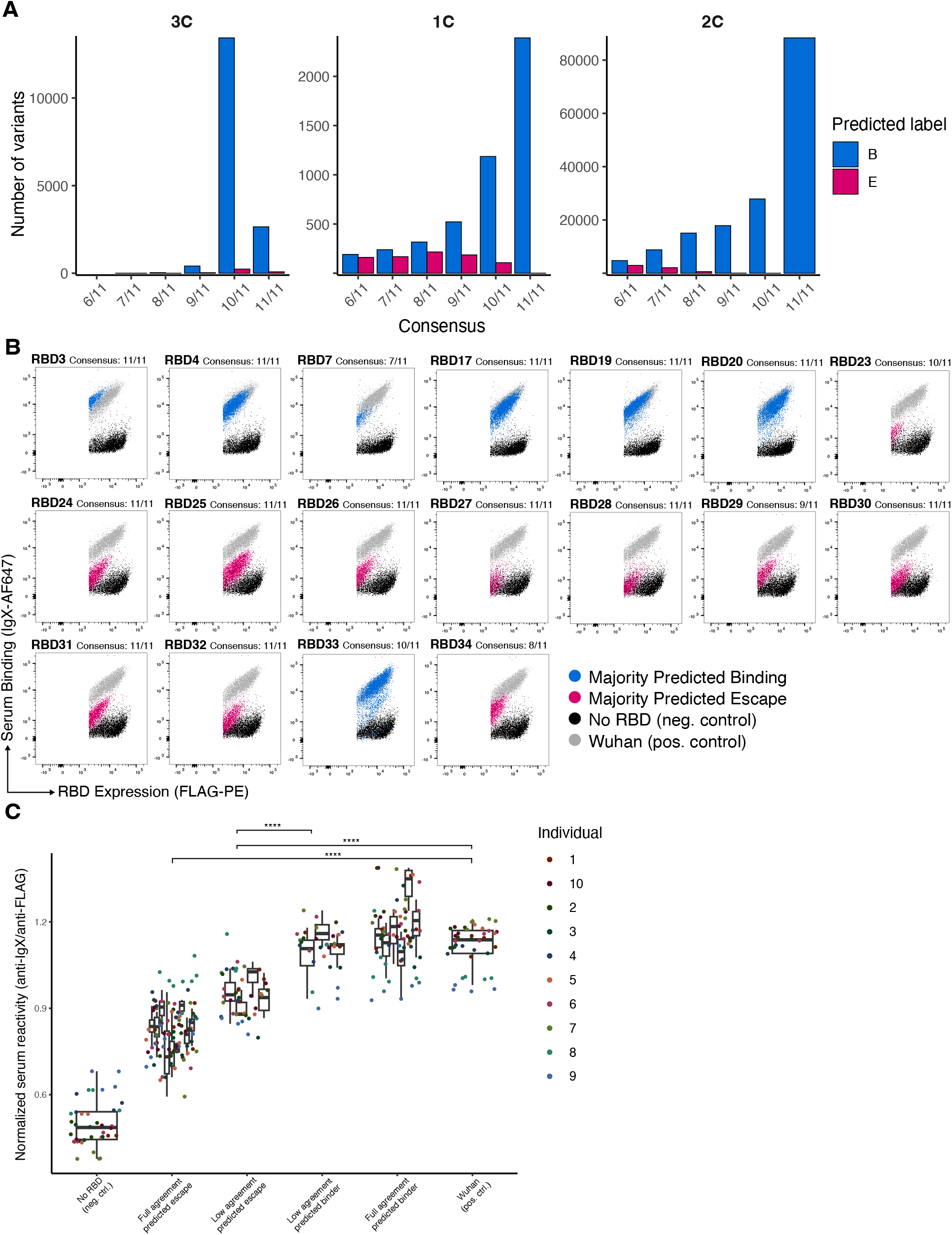
Consensus of trained models is consistent with experimental validations. A) Bar plots showing the degree of agreement between trained classifiers on unseen RBD variants (not present in training or test data). B) Rest of representative FACS plots from screening serum against unseen RBD variants, where the ensemble of all models had full agreement. C) Group comparison of normalized serum reactivity (mean signal ratio anti-IgX/anti-Flag ratios (anti-IgX/anti-Flag) for observed serum binding and -escape variants for all 8 tested observed variants. Statistical analysis was performed across groups using the unpaired two-tailed Student’s t test. ∗p<0.05, ∗∗p<0.01, ∗∗∗p<0.001, ∗∗∗∗p<0.0001, not significant (ns).

**Figure S6.**
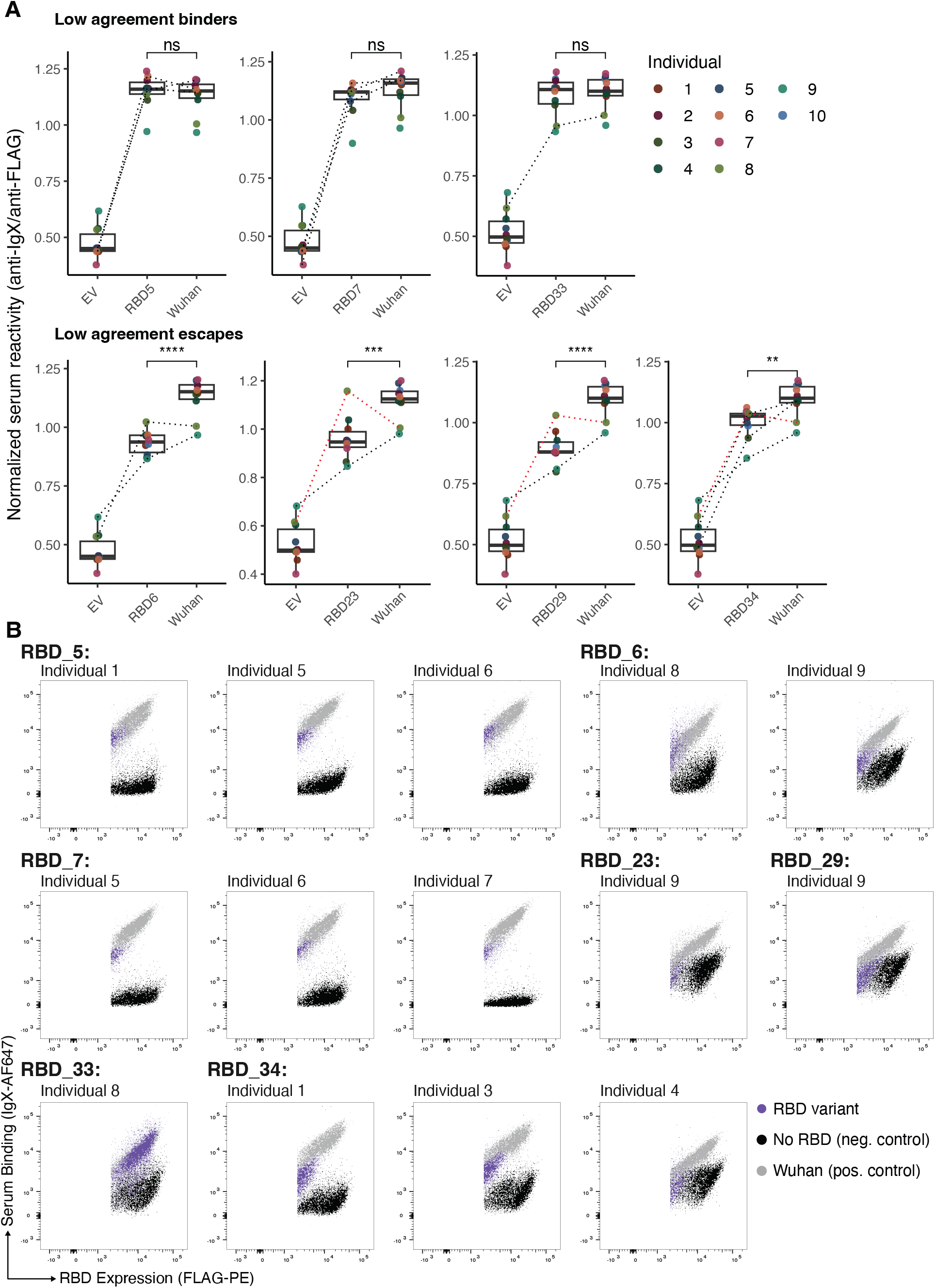
Majority of low agreement predictions are consistent with experimental validations. A) Comparison of normalized serum reactivity (anti-IgX/anti-Flag) for predicted serum binding- and escape-variants with low ensemble model agreement. Black dotted lines represent minority predictions and red dotted lines represent majority predictions that defy the trend. B) FACS plots from screening serum against RBD variants. The shown individuals and RBD variants represent the disagreement of the minority of the consensus predictions. Dotted lines represent minority predictions or majority predictions that are incorrect for the specific individual. Statistical analysis was performed using the unpaired two-tailed Student’s t test. ∗p<0.05, ∗∗p<0.01, ∗∗∗p<0.001, ∗∗∗∗p<0.0001, not significant (ns).

**Table S3.**
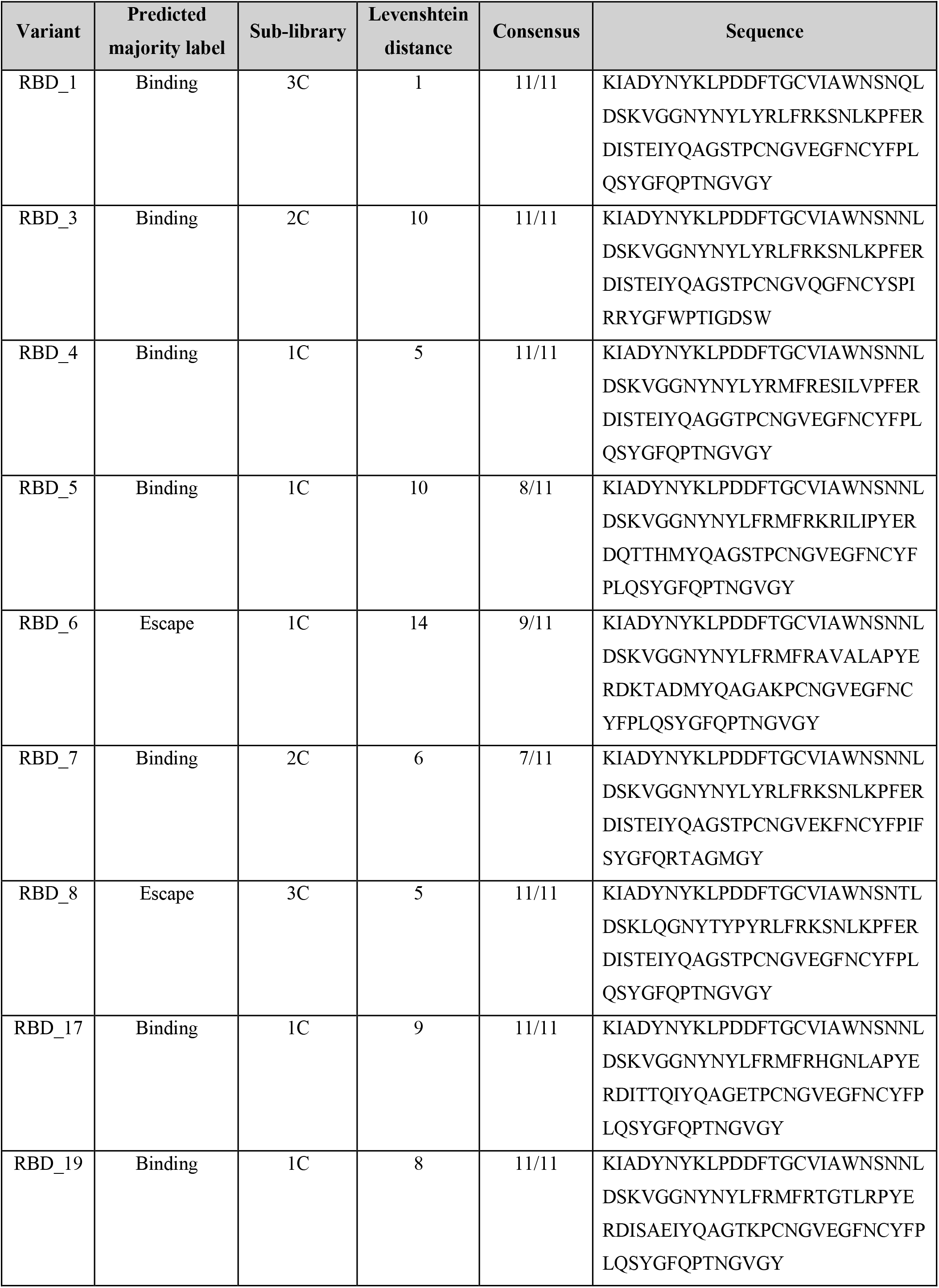
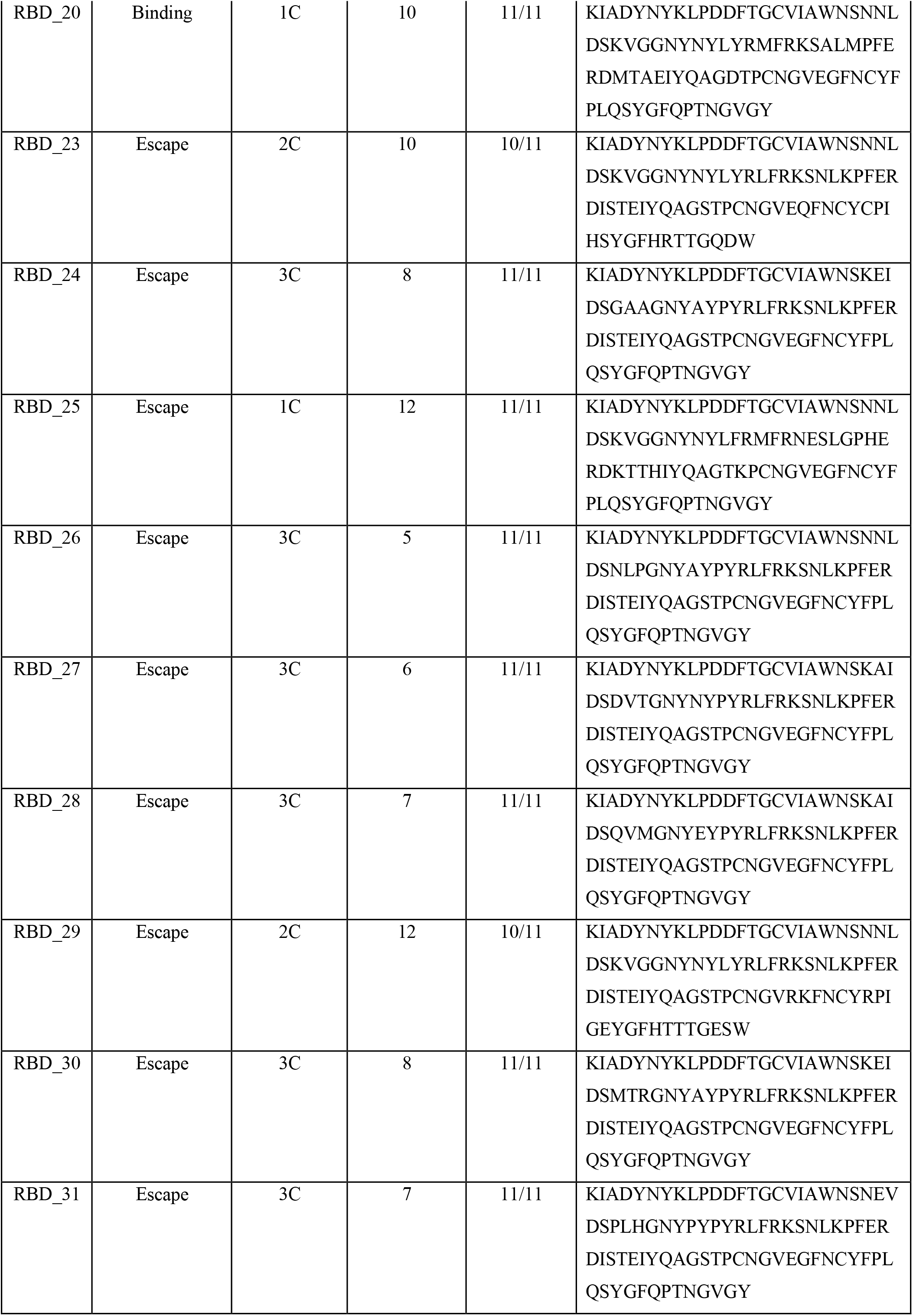
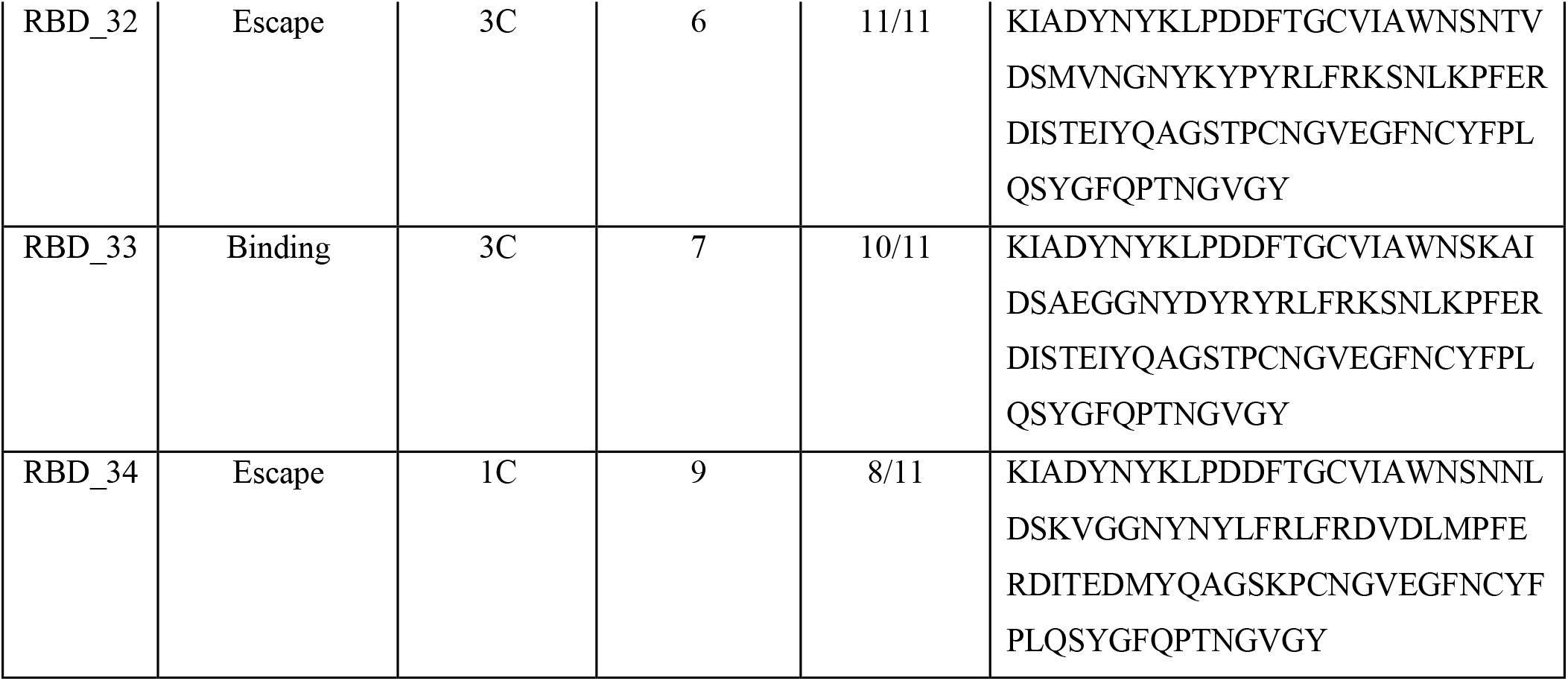
Overview of validated unobserved RBD variants. Variant name, Predicted label (majority of consensus), sub-library to which it belongs, Levenshtein distance from Wu-Hu-1 sequence and consensus of models.

**Table S4.** Overview of low agreement predicted variants. Variant name, Predicted label for each model and Majority prediction.

| Model \ Variant | RBD_33 | RBD_34 | RBD_6 | RBD_5 | RBD_7 | RBD_23 | RBD_29 |
| --- | --- | --- | --- | --- | --- | --- | --- |
| **Cohort-wide** | Binding | Escape | Escape | Binding | Escape | Escape | Escape |
| **Individual 1** | Binding | Escape | Escape | Escape | Binding | Escape | Escape |
| **Individual 2** | Binding | Escape | Escape | Binding | Binding | Escape | Escape |
| **Individual 3** | Binding | Binding | Escape | Binding | Binding | Escape | Escape |
| **Individual 4** | Binding | Binding | Escape | Binding | Binding | Escape | Escape |
| **Individual 5** | Binding | Escape | Escape | Escape | Escape | Escape | Escape |
| **Individual 6** | Binding | Escape | Escape | Escape | Escape | Escape | Escape |
| **Individual 7** | Binding | Escape | Escape | Binding | Escape | Escape | Escape |
| **Individual 8** | Escape | Escape | Binding | Binding | Binding | Escape | Escape |
| **Individual 9** | Binding | Binding | Binding | Binding | Binding | Binding | Binding |
| **Majority Prediction** | Binding | Escape | Escape | Binding | Binding | Escape | Escape |

**Table S5.** Key metrics for Logistic regression, Random forest and RNN models. Parameter tuning for the machine learning models was performed using a grid search approach, using custom scripts and optimizing model performance based on MCC.

| Model | Tuning parameters | Best parameters |
| --- | --- | --- |
| Logistic Regression | max_iter: [1000, 2000, 3000, 4000]<br>C: [1, 10, 50]<br>Solver: [lbfgs, saga] | max_iter: 1000<br>C: 10<br>Solver: lbfgs |
| Random Forest | n_estimators: [400, 500, 550]<br>min_samples_split: [2, 5]<br>min_samples_leaf: [1, 2]<br>max_depth: [100, 150, 200]<br>max_features: [sqrt, None] | n_estimators: 500<br>min_samples_split: 2<br>min_samples_leaf: 1<br>max_depth: 200<br>max_features: sqrt |
| RNN | batch_size: [16, 32]<br>drop_out: [0.1, 0.2]<br>optimizer: [adam, SGD]<br>units: [40, 80]<br>learn_rate: [5e-4, 1e-4, 1e-3] | batch_size: 32<br>drop_out: .2<br>optimizer: adam<br>units: 80<br>learn_rate: 1e-3<br>max epochs: 250 with early stopping<br>patience: 10 |
| Transformer | Tuning was performed based on the default PLMFit parameters:<br>Learning rate: [1e-2 to 1e-6]<br>Batch size: [16 to 512]<br>Weight decay: [1e-4 to 1e-6] | Best parameters were chosen for each run based on smallest validation loss, according to the default PLMfit fine tuning parameters. |

## References

Abebe, E.C. & Dejenie, T.A., 2023. Protective roles and protective mechanisms of neutralizing antibodies against SARS-CoV-2 infection and their potential clinical implications. Frontiers in immunology, 14, p.1055457.

Amanat, F. et al., 2021. SARS-CoV-2 mRNA vaccination induces functionally diverse antibodies to NTD, RBD, and S2. Cell, 184(15), pp.3936–3948.e10.

Barnes, C.O. et al., 2020. Structures of human antibodies bound to SARS-CoV-2 spike reveal common epitopes and recurrent features of antibodies. Cell, 182(4), pp.828–842.e16.

Bikias, T., Stamkopoulos, E. & Reddy, S.T., 2025. PLMFit : Benchmarking transfer learning with protein language models for protein engineering. bioRxiv, p.2025.01.15.633186. Available at: https://www.biorxiv.org/content/10.1101/2025.01.15.633186v1.abstract [Accessed February 10, 2025].

Boder, E.T. & Wittrup, K.D., 1997. Yeast surface display for screening combinatorial polypeptide libraries. Nature biotechnology, 15(6), pp.553–557.

Cao, Y. et al., 2023. Imprinted SARS-CoV-2 humoral immunity induces convergent Omicron RBD evolution. Nature, 614(7948), pp.521–529.

Cao, Y. et al., 2022. Omicron escapes the majority of existing SARS-CoV-2 neutralizing antibodies. Nature, 602(7898), pp.657–663.

Chen, E.C. et al., 2021. Convergent antibody responses to the SARS-CoV-2 spike protein in convalescent and vaccinated individuals. Cell reports, 36(8), p.109604.

Dadonaite, B. et al., 2025. SARS-CoV-2 neutralizing antibody specificities differ dramatically between recently infected infants and immune-imprinted individuals. bioRxivorg, p.2025.01.17.633612. Available at: https://www.biorxiv.org/content/10.1101/2025.01.17.633612v1.abstract [Accessed February 14, 2025].

Dejnirattisai, W. et al., 2021. The antigenic anatomy of SARS-CoV-2 receptor binding domain. Cell, 184(8), pp.2183– 2200.e22.

Eguia, R.T. et al., 2021. A human coronavirus evolves antigenically to escape antibody immunity. PLoS pathogens, 17(4), p.e1009453.

Ehling, R.A. et al., 2024. Synthetic coevolution reveals adaptive mutational trajectories of neutralizing antibodies and SARS-CoV-2. bioRxiv, p.2024.03.28.587189. Available at: https://www.biorxiv.org/content/10.1101/2024.03.28.587189 [Accessed April 15, 2024].

Fowler, D.M. & Fields, S., 2014. Deep mutational scanning: a new style of protein science. Nature methods, 11(8), pp.801–807.

Frei, L. et al., 2023. Deep learning-guided selection of antibody therapies with enhanced resistance to current and prospective SARS-CoV-2 Omicron variants. Immunology. Available at: https://www.biorxiv.org/content/10.1101/2023.10.09.561492v1.full.pdf.

Frei, L. et al., 2025. Deep mutational learning for the selection of therapeutic antibodies resistant to the evolution of Omicron variants of SARS-CoV-2. Nature biomedical engineering, 9(4), pp.552–565.

Glick, M. et al., 2003. Prioritization of high throughput screening data of compound mixtures using molecular similarity. Molecular physics, 101(9), pp.1325–1328.

Greaney, A.J., Starr, T.N., Gilchuk, P., et al., 2021. Complete mapping of mutations to the SARS-CoV-2 spike receptor-binding domain that escape antibody recognition. Cell host & microbe, 29(1), pp.44–57.e9.

Greaney, A.J., Loes, A.N., et al., 2021. Comprehensive mapping of mutations in the SARS-CoV-2 receptor-binding domain that affect recognition by polyclonal human plasma antibodies. Cell host & microbe, 29(3), pp.463– 476.e6.

Greaney, A.J., Starr, T.N., Barnes, C.O., et al., 2021. Mapping mutations to the SARS-CoV-2 RBD that escape binding by different classes of antibodies. Nature communications, 12(1), p.4196.

Greaney, A.J., Starr, T.N. & Bloom, J.D., 2021. An antibody-escape calculator for mutations to the SARS-CoV-2 receptor-binding domain. bioRxivorg. Available at: https://jbloomlab.github.io/RBD_escape_calculator_paper/paper.pdf.

Lee, J.M. et al., 2019. Mapping person-to-person variation in viral mutations that escape polyclonal serum targeting influenza hemagglutinin. eLife, 8. Available at: https://elifesciences.org/articles/49324 [Accessed February 10, 2025].

Lima, N.S. et al., 2022. Primary exposure to SARS-CoV-2 variants elicits convergent epitope specificities, immunoglobulin V gene usage and public B cell clones. Nature communications, 13(1), p.7733.

Lopez-Morales, J. et al., 2023. Multiplexed on-yeast serological assay for immune escape screening of SARS-CoV-2 variants. iScience, 26(5), p.106648.

Ma, W. et al., 2023. Immune evasion and ACE2 binding affinity contribute to SARS-CoV-2 evolution. Nature ecology & evolution, 7(9), pp.1457–1466.

Minot, M. & Reddy, S.T., 2024. Meta learning addresses noisy and under-labeled data in machine learning-guided antibody engineering. Cell systems, 15(1), pp.4–18.e4.

Piccoli, L. et al., 2020. Mapping Neutralizing and Immunodominant Sites on the SARS-CoV-2 Spike Receptor-Binding Domain by Structure-Guided High-Resolution Serology. Cell, 183(4), pp.1024–1042.e21.

Starr, T.N., Greaney, A.J., Dingens, A.S., et al., 2021. Complete map of SARS-CoV-2 RBD mutations that escape the monoclonal antibody LY-CoV555 and its cocktail with LY-CoV016. Cell reports. Medicine, 2(4), p.100255.

Starr, T.N. et al., 2020. Deep Mutational Scanning of SARS-CoV-2 Receptor Binding Domain Reveals Constraints on Folding and ACE2 Binding. Cell, 182(5), pp.1295–1310.e20.

Starr, T.N., Greaney, A.J., Addetia, A., et al., 2021. Prospective mapping of viral mutations that escape antibodies used to treat COVID-19. Science (New York, N.Y.), 371(6531), pp.850–854.

Starr, T.N., Czudnochowski, N., et al., 2021. SARS-CoV-2 RBD antibodies that maximize breadth and resistance to escape. Nature, 597(7874), pp.97–102.

Starr, T.N. et al., 2022. Shifting mutational constraints in the SARS-CoV-2 receptor-binding domain during viral evolution. Science (New York, N.Y.), 377(6604), pp.420–424.

Taft, J.M. et al., 2022. Deep mutational learning predicts ACE2 binding and antibody escape to combinatorial mutations in the SARS-CoV-2 receptor-binding domain. Cell, 185(21), pp.4008–4022.e14.

Tao, K. et al., 2021. The biological and clinical significance of emerging SARS-CoV-2 variants. Nature reviews. Genetics, 22(12), pp.757–773.

Traxlmayr, M.W., 2022. Yeast Surface Display.

Yan, Q. et al., 2024. Antibodies utilizing VL6-57 light chains target a convergent cryptic epitope on SARS-CoV-2 spike protein and potentially drive the genesis of Omicron variants. Nature communications, 15(1), p.7585.

Yuan, M. et al., 2021. Structural and functional ramifications of antigenic drift in recent SARS-CoV-2 variants. Science (New York, N.Y.), 373(6556), pp.818–823.

Zhang, A. et al., 2023. Beyond neutralization: Fc-dependent antibody effector functions in SARS-CoV-2 infection. Nature reviews. Immunology, 23(6), pp.381–396.

